# How benthic sediment microbial communities respond to glyphosate and its metabolite: A microcosm experiment

**DOI:** 10.1101/2022.12.30.522317

**Authors:** Christine M Cornish, Peter Bergholz, Kaycie Schmidt, Jon Sweetman

**Author notes:** Corresponding author: Christine Cornish.

## Abstract

Glyphosate is the most commonly used agricultural herbicide in the world. In aquatic ecosystems, glyphosate often adsorbs to benthic substrates or is metabolized and degraded by microorganisms. The effects of glyphosate on microbial communities varies widely as microorganisms respond differently to exposure. To help understand the impacts of glyphosate on the sediment microbiome we conducted a microcosm experiment examining the responses of benthic sediment microbial communities to herbicide treatments. Sediments from a prairie pothole wetland were collected and 16S rRNA gene sequencing was used to analyze community composition 2-hours and 14-days after a single treatment of low (0.07 ppm), medium (0.7 ppm), or high (7 ppm) glyphosate, aminomethylphosphonic acid (glyphosate metabolite), or a glyphosate-based commercial formula. We found no significant differences in microbial community composition between treatments, concentration levels, or time. These findings suggest that microbial species in the Prairie Pothole Region of North America may be tolerant to glyphosate exposure.

## Introduction

Agrochemical contamination of aquatic ecosystems is an ongoing concern due to the direct and indirect risks to environmental health. Glyphosate (i.e., Roundup^®^) is a non-selective, systemic herbicide that has become the most commonly used herbicide in the world since the 1990s [1]. The substantial use of glyphosate has resulted in its widespread and frequent detection in surface waters and groundwater [2, 3], where benthic sediments often become sinks [4]. Microbial metabolism is the primary degradation mechanism of glyphosate [5] resulting in its metabolites, aminomethylphosphonic acid (AMPA) or phosphate and sarcosine [6].

Glyphosate targets higher plants through the shikimate pathway [7], but this pathway is also present in some microorganisms [8]. Therefore, it was initially presumed that glyphosate may have inadvertent effects on microbial communities. Studies have reported that glyphosate can have a wide range of negative, positive, or neutral impacts on microbial community composition and function in terrestrial and aquatic ecosystems. Aquatic microbial communities have been shown to utilize glyphosate as a nutrient source resulting in increased activity, whereas in others, microbes are inhibited by toxicological effects. Pérez et al. [9] found 6 mg L^-1^ and 12 mg L^-1^ Roundup^®^ caused a significant decrease in average microphytoplankton and nanophytoplankton abundance, whereas Lu et al. [10] found significantly enhanced gene expression in mechanisms potentially related to glyphosate tolerance, but did not find overall community structure shifts from 2.5 mg L^-1^ glyphosate. Several species are known to exhibit glyphosate tolerance [6], in addition to species capable of using glyphosate directly as a nutrient source [11, 12]. These examples demonstrate the variety of impacts glyphosate can have on aquatic microbial communities.

In addition to the impacts of glyphosate, AMPA, glyphosate’s main metabolite, can also have direct and indirect effects on aquatic ecosystems [13]. Similar to glyphosate AMPA can be degraded by microorganisms, but it is more mobile and persistent [14–16]. It is also a known phytotoxin [17] with additional concerns regarding its potential to bioaccumulate [18]. AMPA is highly dependent on the presence and concentration of glyphosate [19], where both compounds frequently co-occur in areas of high agricultural intensity [5, 20]. Glyphosate and AMPA are most often detected in surface soil [21], and frequently transported to surface waters, including wetlands [2, 5] where they can impact benthic microorganisms.

The Prairie Pothole Region (PPR) is a large complex landscape covered with shallow wetlands and prairies [22]. This region is surrounded by agriculture, primarily corn and soybean croplands, which are the predominant crops glyphosate is used on [1, 2]. Consequently, wetlands in the PPR are subject to prolonged glyphosate contamination, where glyphosate has been reported as the most frequently detected and the highest detected herbicide in this region [23]. These wetlands are key ecosystems providing many economical and ecological services. For example, biogeochemical processes including C turnover and sequestration, N and P capture, and remediation of agrochemicals are essential ecosystem functions [24, 25], where benthic sediment microbial communities play a fundamental role in these ecosystem processes.

Understanding the impacts of agrochemical contamination on microbial community structure is vital because shifts in microbial communities impact the whole ecosystem. This is especially important in a region with high agricultural intensity, like the PPR, because microorganisms are chronically exposed to chemical stressors like glyphosate. The objective of our research was to assess the responses of benthic sediment microbial communities from the PPR to glyphosate-based herbicide treatments. We hypothesized that 1) observed Operational Taxonomic Units (OTUs) and microbial diversity would decrease at high herbicide concentration due to toxicological effects, and 2) microbial community composition would shift at low herbicide concentration in favor of species capable of metabolizing glyphosate.

## Materials and Methods

### Microcosm preparation

Surface sediment (∼ 10 cm) was collected from a wetland (P1) within the Cottonwood Lake Study Area (U.S. Fish and Wildlife service managed Waterfowl Production Area) of the PPR located in Stutsman County, North Dakota. This area has minimal agricultural influence, where native prairie grasslands and wetlands cover over 80% [26]. After collection, sediment was stored refrigerated (∼ 4 °C) until initiation of the experiment. Thirty-two 5.7 L microcosms were prepared with the following contents: 1 cm layer of homogenized sediment, 2.5 L of dechlorinated tap water, and covered with “no-see-um” mesh (Duluth Sport Nets, Duluth, MN). Microcosms were then stored in incubators at 20 °C with a 16:8 (light:dark) hour photoperiod and left to acclimate for one month prior to treatment. Over the entirety of the 6-week experiment, microcosms were monitored for water evaporation and were filled back to the 2.5 L volume when necessary.

### Herbicide treatment and sediment sampling

We conducted a 3 x 4 factorial experiment with three replicates per treatment and five controls (*n* = 32). Herbicide treatments consisted of analytical grade glyphosate (98.1% purity) or AMPA (99% purity), or 41% glyphosate concentrate (commercial formula) at concentrations of 0.07 parts per million (ppm), 0.7 ppm, or 7 ppm. Our concentrations were chosen based on the U.S. maximum contaminant level (MCL) of glyphosate in drinking water (U.S. EPA 2015), which is equivalent to our medium concentration of 0.7 ppm (or 7 mg L^-1^). Treatments were added to the water, lightly stirred to evenly distribute, and then allowed to settle for two hours. A 50 mL polypropylene corer (needle-less syringe) was used to collect sediment samples pretreatment, two hours post-treatment, and two weeks post-treatment. All samples were stored in 50 mL polypropylene tubes at -80 °C immediately after sampling until analyses were performed.

### Analyses: Herbicide residues and microbial 16S sequencing

Sediment subsamples were shipped frozen on dry ice to the Agriculture and Food Laboratory at the University of Guelph (Ontario, Canada) for glyphosate and AMPA residue analysis. Liquid chromatography tandem mass spectrometry (LC-MS/MS) was used to quantify glyphosate and AMPA residues with a limit of detection of 0.005 ppm and a limit of quantification of 0.02 ppm for both compounds. Matrix blanks and standards were analyzed alongside sediment samples for quality control (QC).

Sediment microbial communities were analyzed after extraction of environmental DNA with the Qiagen DNeasy^®^ PowerSoil^®^ kit. Briefly, 0.25 g sediment (wet weight) was lysed using PowerBead Tubes in a bead-beater (Biospec Mini-beadbeater-24). Kit solutions (C1 – C6) were added stepwise to purify, wash, and elute DNA into 100 μL final volume. Microbial 16S rRNA gene was PCR amplified using universal primers, 27F and U1492R, for QC to verify suitability for sequencing. Sequencing library preparation and sequencing were performed according to Oxford Nanopore Technologies 16S Barcoding Kit (SQK-RAB204) protocol and reagents. Briefly, amplicons were cleaned using AMPure XP bead cleanup, quantified via PicoGreen™ analysis (Quant-iT PicoGreen™ Kit, modified from Invitrogen’s Quant-iT PicoGreen™ dsDNA Reagent and Kits protocol), and pooled to obtain a final DNA concentration of 50 – 100 ng. Pooled samples contained up to twelve separately barcoded samples for sequencing using an Oxford Nanopore minION™. Each library was run for approximately four hours. Raw fast5 reads were basecalled and demultiplexed using Guppy v3.4. EPI2ME 16S workflow (https://epi2me.nanoporetech.com, rev 2.1.0) was used for QC and initial characterization. For QIIME2 analysis, sequences were pre-processed using MetONTIIME [27] and QIIME2 was used to filter and analyze the resulting OTU table abundances. The SILVA v138 database was downloaded and utilized as a reference database for taxonomic identification [28, 29]. Taxonomy barplots, describing the composition of each sample at the desired taxonomic level were visualized at QIIME2 view (https://view.qiime2.org/). Sequence tables were filtered to remove taxonomically unassigned sequences. A table of absolute OTU abundances was exported in BIOM format for further analysis in R.

### Statistical analysis

A feature-table containing absolute abundances of Family level OTUs from QIIME2 was imported into R (version 4.0.2) using the R/biomformat package (1.24.0) [30]. Unassigned taxa and taxa that were present in ≤ 10% of samples were removed to minimize the risk of sequencing error carryover and due to low statistical power to analyze underrepresented OTUs, respectively. All subsequent analyses were conducted using the R/stats (3.6.2) and R/vegan (2.6-2) packages [31, 32]. Shannon’s H. diversity index was calculated using the “diversity” function and observed OTUs were calculated using the “specnumber” function. Differences in diversity and observed OTUs between herbicide treatments, concentration levels, and time of sampling were analyzed with a Kruskal-Wallis Rank Sum test using the “kruskal.test” function. Lastly, Bray-Curtis dissimilarity distance measures were calculated and square root transformed for non-metric multidimensional scaling (NMDS) using the “metaMDS” function to display community structure.

## Results

After unassigned and rare taxa were removed, a total of 430 OTUs were detected across all sediment samples (*n* = 96). Relative abundances across treatments were not significantly different (Fig. 1). The three most abundant families in all treatments were *Hydrogenophilaceae* in the *Pseudomonadota* (*Gammaproteobacteria*) Phylum, and *Sulfurovaceae* and *Desulfobacteraceae*, which are assigned to the *Proteobacteria* (*Deltaproteobacteria*) Phylum. *Prolixibacteraceae*, a family within the *Bacteroidota* Phylum thought to be associated with sediment, was only detected in the control and glyphosate treatments.

**Fig. 1.**
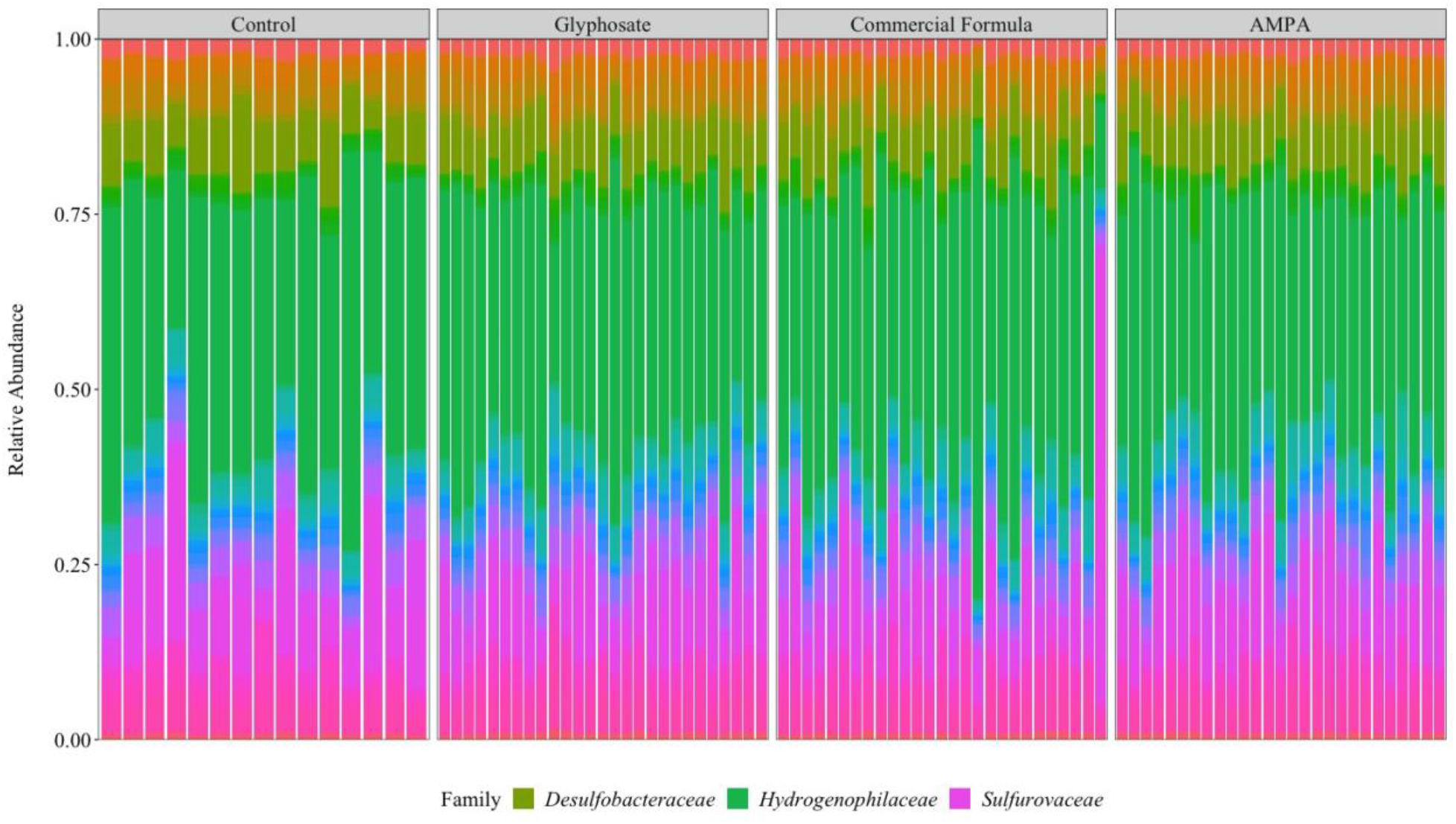
Relative abundance of microbial Families among all samples (*n* = 96) in each herbicide treatment; legend only shows the top three most abundant Families across all herbicide treatments

Shannon H index and observed OTUs were similar among all treatments and controls over the entirety of the experiment (Fig. 2 and Fig. 3). Kruskal-Wallis tests showed no significant differences in alpha diversity metrics between herbicide treatments, concentration levels, or time of sampling (Table 1). A two-dimensional ordination solution was reached (stress = 0.1793708) using NMDS. Microbial communities were similar in multivariate space, which showed there were no significant compositional differences between treatments (Fig. 4).

**Table 1.** Summary of Kruskal-Wallis Rank Sum test on the effects of treatments on Shannon diversity and observed OTU. *χ*^2^ = Chi-square value, df = degrees of freedom, and *P* = the p-value where the significance level is 0.05

| Quantified Metric | Treatment | $\chi^2$ | df | $P$ |
| --- | --- | --- | --- | --- |
| Shannon Diversity | Time of Sampling | 1.81 | 2 | 0.40 |
|  | Herbicide Treatment | 4.67 | 3 | 0.20 |
|  | Concentration | 3.48 | 3 | 0.32 |
| Observed OTU | Time of Sampling | 2.25 | 3 | 0.52 |
|  | Herbicide Treatment | 0.22 | 2 | 0.90 |
|  | Concentration | 2.45 | 3 | 0.48 |

**Fig. 2.**
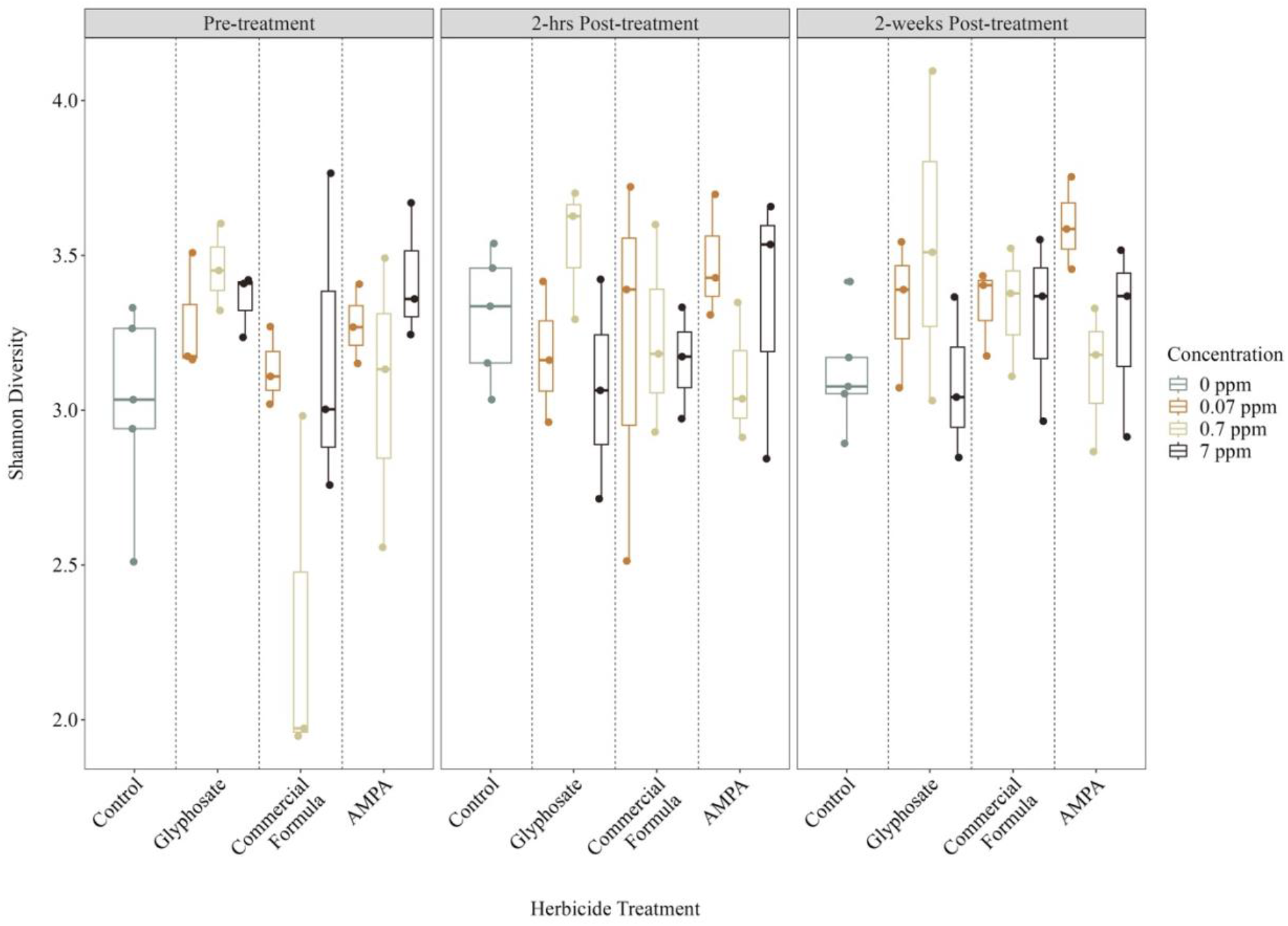
Shannon H diversity index quantified from microcosm sediments of herbicide treatments and treatment levels at each sampling timepoint

**Fig. 3.**
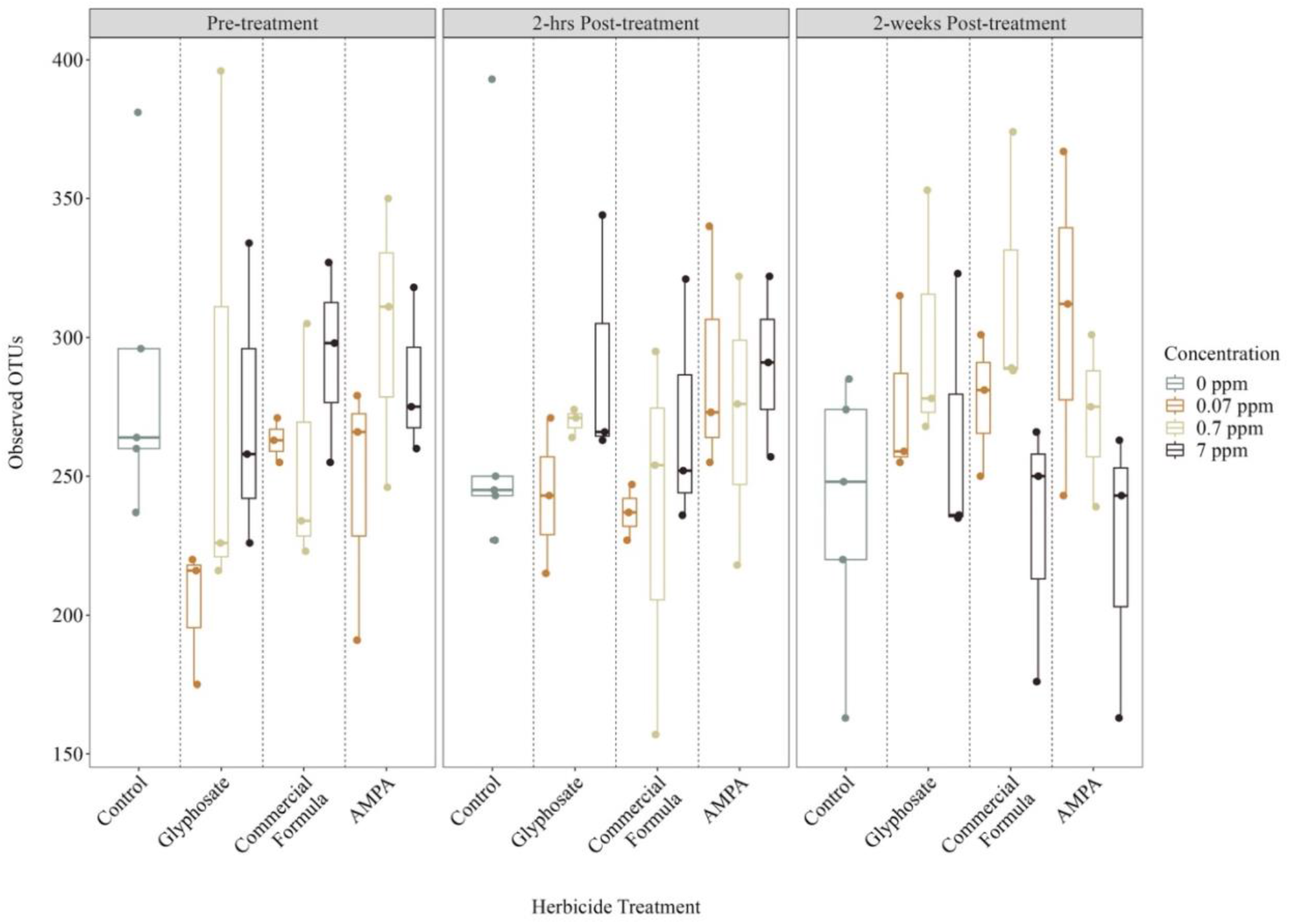
Observed OTUs quantified from microcosm sediments of herbicide treatments and treatment levels at each sampling timepoint

**Fig. 4.**
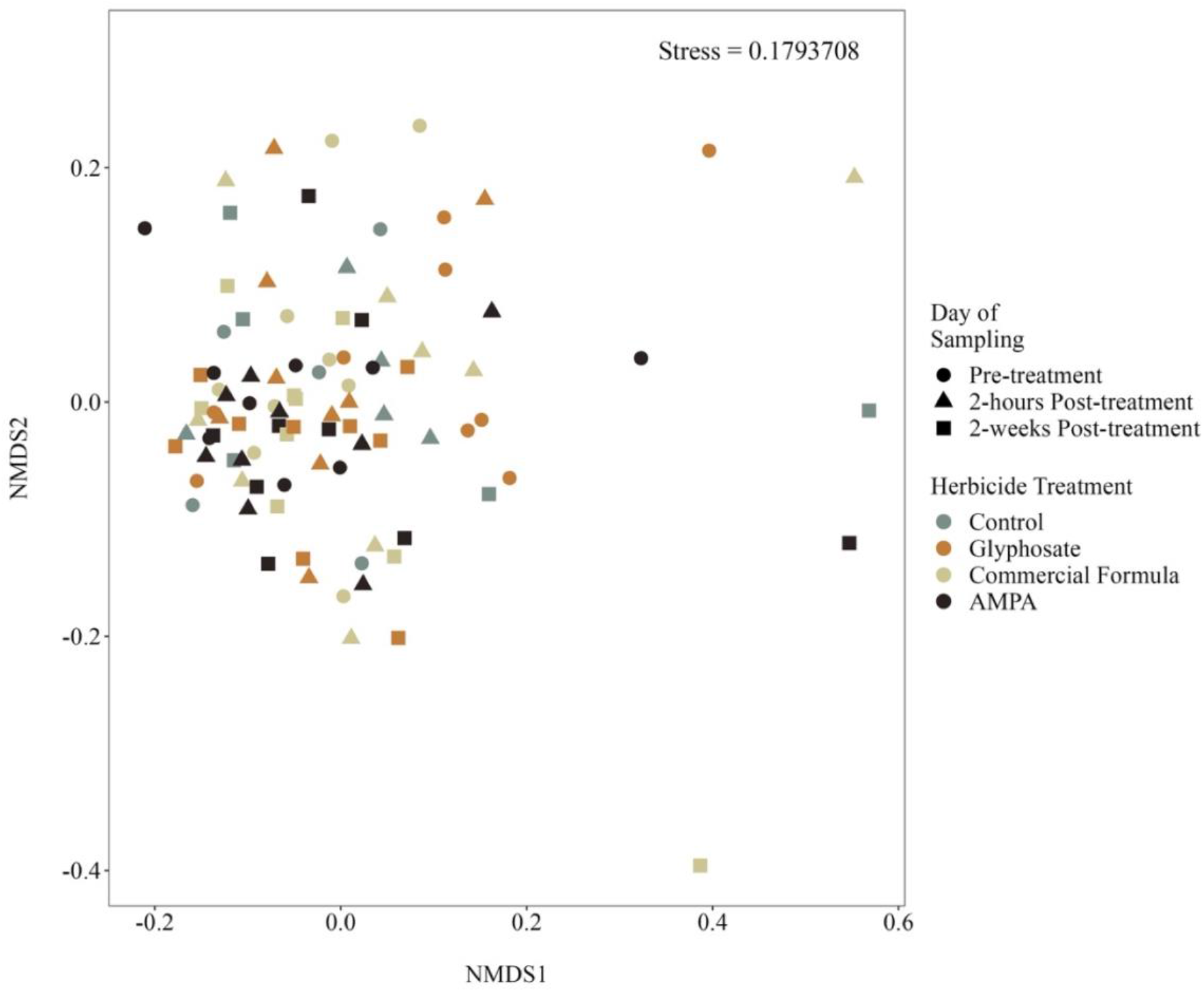
NMDS plot of sediment microbial community structure across all samples (*n* = 96) where shapes represent day of sampling and colors represent herbicide treatment

Both glyphosate and AMPA concentrations in sediments increased over time within each herbicide treatment, but not in control treatments. Additionally, AMPA was detected in microcosms where glyphosate or commercial formula only was added, confirming that glyphosate degradation did occur (Online Resource 1).

## Discussion

The current study used microcosms to examine the impact of glyphosate-based herbicide treatments on benthic sediment microbial communities from a prairie pothole wetland. Both glyphosate and AMPA were detected in microcosm sediments shortly following treatment indicating that residues rapidly dissipate out of the water column [5, 15]. AMPA was also detected in sediments of microcosms that only received glyphosate (i.e., glyphosate or commercial formula treatment), but no direct AMPA treatment, indicating that glyphosate degradation occurred [33]. In our experimental microcosms, we found that microbial community diversity and composition were not significantly affected by glyphosate, AMPA, or commercial formula at any concentration, which was contrary to our original hypotheses. Our highest treatment level (7 ppm) was a magnitude higher than the U.S. EPA’s MCL for drinking water, and approximately 41 times higher than the average concentration that have been reported in wetland sediments in the PPR [23]. Some research suggests that microbial responses are more dependent on previous glyphosate exposure history and application rates [18, 34]. The present experiment reflects conditions of acute glyphosate and AMPA exposure, rather than long-term exposure on the sediment community. Sediments used in our study were collected from a permanent wetland in North Dakota with no routine pesticide usage in the immediate catchment. Thus, agricultural inputs within this area are minimal to none compared to other areas of the PPR. However, we found no differences between treatments and controls suggesting that glyphosate-based herbicides may not have adverse effects on sediment microbial communities from wetlands in this area of the PPR. This observed lacked effect may be the result of glyphosate- and AMPA-based selection pressure.

A lack of a response from microbial communities following glyphosate exposure has also been observed previously in the literature. For example, Pesce et al. [35] exposed natural spring- and summer-collected riverine microbial communities to 10 μg L^-1^ glyphosate and found no effect on bacterial activity or diversity. Lane et al. [36] conducted a six-month soil incubation experiment with monthly glyphosate treatments at 59 μg g^-1^ and 118 μg g^-1^, and found no significant glyphosate effect on community structure represented by the relative abundances of functional microbial groups. Muturi et al. [37] used 20 mg L^-1^ glyphosate treatment in microcosms and found no differences in water microbial diversity or richness after 3- and 7-days. Dennis et al. [38] found no significant effects of a single glyphosate treatment at the recommended field application rate (33.03 mg L^-1^) on bacterial or archaeal evenness, richness, and composition after 60-days incubation. These studies in addition to the current study represented a wide range of glyphosate concentrations, which all showed that glyphosate does not always have direct toxic effects on microbial communities. Presumably, this may be due to the presence of glyphosate-tolerant microbes.

Several studies have, however, shown shifts in microbial communities after either short- or long-term glyphosate exposure. Lu et al. [10], for example, found increases in Shannon and Simpson diversity of lacustrine microbial communities, in addition to differences in community structure 10- and 15-days post-treatment of 2.5 mg L^-1^ glyphosate. Widenfalk et al. [39] also found that an environmentally relevant glyphosate concentration (150 μg kg^-1^ dry weight) caused significant shifts in lake sediment bacterial community composition in treated microcosms relative to controls after 31-days. Sura et al. [40] found pelagic and biofilm bacterial production in outdoor mesocosms was significantly inhibited by 225 μg L^-1^ glyphosate compared to controls. Microbial communities in our study and Sura et al. [40] were both collected from wetlands within the PPR, however our results were not consistent potentially due to differences in sediment-versus water-associated communities, land use within our collection site’s watersheds, or incubation versus outdoor experimental design. These studies all showed that glyphosate can have various negative and positive effects on microbial communities, whereas our results showed glyphosate can also have no effect. This discrepancy indicates that there are many underlying complexities on the effects of glyphosate at the microbial level, which may be more related to molecular mechanisms and/or environmental variables.

Some microorganisms naturally express a glyphosate-resistant form of the EPSP enzyme and there are many glyphosate-tolerant microbial species listed in the literature [41, 42]. Specifically, *Agrobacterium* sp. strain CP4 was the bacteria used for the original glyphosate-resistant EPSP gene in glyphosate-resistant crops [43]. Whereas, other species have evolved tolerance through mutations of the EPSP synthase, or metabolic or detoxifying processes [44], which can be a result of prolonged or repeated glyphosate exposure. For example, Tang et al. [34] added 100 mg L^-1^ glyphosate to sediments with high, low, and no previous glyphosate exposure, and found that microbes degraded glyphosate quicker in sediments with high exposure history compared to low exposure history. Additionally, Lane et al. [36] found significantly higher microbial respiration in soils after glyphosate treatment, specifically in soils with previous glyphosate exposure history. The sediments used in the present study were collected from an agriculturally-undisturbed wetland located in an area with no known extensive glyphosate use (David Mushet, Research Wildlife Biologist, USGS Northern Prairie Wildlife Research Center, email communication June 21, 2022), thus microbial communities should not have any prolonged exposure history. We did find multiple species across all treatments that are capable of glyphosate degradation and mineralization including *Cyanobiaceae, Enterobacteriaceae, Pseudomonadaceae*, and *Rhizobiaceae* (e.g., genus *Agrobacterium*). While our sediment collection site has no known history of glyphosate use within the immediate watershed, the majority of the PPR is in an agriculturally intensive area where glyphosate use is common. Therefore, microbial species from the regional species pool may have evolved tolerance despite our study wetland and its catchment having no existing agricultural pesticide use.

Glyphosate is highly water soluble, which also allows it to be easily transported into wetlands, where environmental variables can play a role in its bioavailability and toxicity to microorganisms. Glyphosate has an affinity for oxides and organic matter content resulting in its rapid adsorption to sediments [45]. However, glyphosate and phosphate can compete for sediment binding sites due to their chemical structural similarities [46]. Thus, lower phosphate can increase glyphosate’s sediment adsorption capacity [47], subsequently decreasing its bioavailability. Temperature can also affect glyphosate’s environmental persistence, where higher temperatures facilitate degradation due to increased microbial activity [48, 49]. While the present study did not measure sediment physicochemical characteristics, temperature and light conditions were controlled for, and homogenized sediments were used for all microcosms to help minimize variability.

### Conclusion

We evaluated the responses of benthic sediment microbial communities to a single addition of low, medium, or high glyphosate-based herbicide treatment after 14-days incubation. We expected to see toxicological-induced shifts potentially in favor of tolerant species, however we found no differences in sediment microbial communities among treatments or concentration after 2 weeks. Our results may be explained by the lower concentration of bioavailable glyphosate in the sediments compared to the actual concentrations added. Additionally, our results may suggest glyphosate-tolerance in the benthic sediment microbial communities, but that is inconclusive without further investigation of the presence of the glyphosate-resistant EPSP gene within the microbiome. Our research suggests that in the PPR the direct effects of glyphosate on sediment microorganisms may not be as severe as initially presumed. However, the literature continues to reveal new implications of the extensive use of glyphosate on aquatic ecosystems. Therefore, further research is still necessary to determine the full range of potential effects of glyphosate on sediment microbial communities.

## Acknowledgments

Thank you to the North Dakota Established Program to Stimulate Competitive Research for funding this research. Thank you to Dr. López-Martínez for providing support and usage of equipment. Additionally, thank you to the anonymous reviewers for their time and feedback of our manuscript.

## Statements and Declarations

### Author contributions

All authors contributed to the study conception and design. Material preparation, data collection, and microbial analysis were performed by Christine Cornish and Kaycie Schmidt. The first draft of the manuscript was written by Christine Cornish with additions from Jon Sweetman and Peter Bergholz. All authors commented on previous versions of the manuscript, and read and approved the final manuscript.

### Availability of data and materials

The data used in this study are available from the corresponding author upon reasonable request.

### Ethics approval

Not applicable.

### Consent to participate

All authors have given consent to participate.

### Consent for publication

All authors have given consent to publish.

