## Supplemental Material for "How benthic sediment microbial communities respond to glyphosate and its metabolite: A microcosm experiment"

**Microbial Ecology**

Christine M Cornish^1^, Peter Bergholz, Kaycie Schmidt, and Jon Sweetman

^1^Department of Biological Sciences, North Dakota State University

1340 Administration Avenue, Fargo, ND 58105, United States

**Supplemental Table 1** Herbicide residue analysis from the Agriculture and Food Laboratory at the University of Guelph (Ontario, Canada). The limit of detection for glyphosate was 0.005 ppm and the limit of quantification (LOQ) was 0.02 ppm. The columns show average glyphosate concentration (ppm/mg L^-1^) ± standard deviation detected in microcosm sediments of each herbicide treatment and concentration at each sampling timepoint. Concentrations: L = 0.07 ppm, M = 0.7 ppm, and H = 7 ppm. If residues were < LOQ it was designated a value of 0.02 for calculation purposes.

|  | | Pre-treatment | 2-hours  Post-treatment | 2-weeks  Post-treatment |
| --- | --- | --- | --- | --- |
| Treatment | Concentration |  |  |  |
| Control |  | 0.01 ± 0.01 | 0.00 ± 0.00 | 0.00 ± 0.00 |
| Glyphosate (purity = 98.1%) | L | 0.03 ± 0.04 | 0.00 ± 0.00 | 0.05 ± 0.08 |
|  | M | 0.00 ± 0.00 | 0.03 ± 0.04 | 0.28 ± 0.1 |
|  | H | 0.00 ± 0.00 | 0.67 ± 0.35 | 4.37 ± 0.35 |
| Commercial Formula | L | 0.00 ± 0.00 | 0.00 ± 0.00 | 0.00 ± 0.00 |
|  | M | 0.00 ± 0.00 | 0.00 ± 0.00 | 0.31 ± 0.36 |
|  | H | 0.00 ± 0.00 | 0.41 ± 0.46 | 2.77 ± 1.30 |
| AMPA  (purity = 99%) | L | 0.00 ± 0.00 | 0.01 ± 0.01 | 0.00 ± 0.00 |
|  | M | 0.00 ± 0.00 | 0.00 ± 0.00 | 0.00 ± 0.00 |
|  | H | 0.01 ± 0.01 | 0.00 ± 0.00 | 0.00 ± 0.00 |

**Supplemental Table 2** Herbicide residue analysis from the Agriculture and Food Laboratory at the University of Guelph (Ontario, Canada). The limit of detection for AMPA was 0.005 ppm and the limit of quantification (LOQ) was 0.02 ppm. The columns show average AMPA concentration (ppm/mg L^-1^) ± standard deviation detected in microcosm sediments of each herbicide treatment and concentration at each sampling timepoint. Concentrations: L = 0.07 ppm, M = 0.7 ppm, and H = 7 ppm. If residues were < LOQ it was designated a value of 0.02 for calculation purposes.

|  | | Pre-treatment | 2-hrs  Post-treatment | 2-weeks  Post-treatment |
| --- | --- | --- | --- | --- |
| Treatment | Concentration |  |  |  |
| Control |  | 0.00 ± 0.00 | 0.00 ± 0.00 | 0.00 ± 0.00 |
| Glyphosate (purity = 98.1%) | L | 0.00 ± 0.00 | 0.00 ± 0.00 | 0.00 ± 0.00 |
|  | M | 0.01 ± 0.01 | 0.01 ± 0.01 | 0.04 ± 0.02 |
|  | H | 0.00 ± 0.00 | 0.05 ± 0.05 | 0.35 ± 0.15 |
| Commercial Formula | L | 0.01 ± 0.01 | 0.00 ± 0.00 | 0.01 ± 0.01 |
|  | M | 0.00 ± 0.00 | 0.00 ± 0.00 | 0.01 ± 0.01 |
|  | H | 0.00 ± 0.00 | 0.03 ± 0.02 | 0.19 ± 0.03 |
| AMPA  (purity = 99%) | L | 0.01 ± 0.01 | 0.01 ± 0.01 | 0.04 ± 0.03 |
|  | M | 0.00 ± 0.00 | 0.11 ± 0.04 | 0.48 ± 0.18 |
|  | H | 0.00 ± 0.00 | 1.08 ± 0.57 | 6.07 ± 7.90 |

**Supplemental Table 3** Average quantified species metrics ± standard deviation detected in microcosm sediments of each herbicide treatment over the entirety of the experiment.

|  | Shannon Diversity | Observed OTUs |
| --- | --- | --- |
| Treatment |  |  |
| Control | 2.7 ± 0.24 | 266 ± 58 |
| Glyphosate  (purity = 98.1%) | 2.9 ± 0.26 | 267 ± 50 |
| Commercial Formula | 2.7 ± 0.41 | 264 ± 45 |
| AMPA  (purity = 99%) | 2.9 ± 0.29 | 276 ± 47 |
